# Harnessing *Escherichia coli* motility to engineer bacterial Voronoi patterns

**DOI:** 10.64898/2026.08.31.748246

**Authors:** Jung Hun Park, Gábor Holló, Emanuele Boni, Yolanda Schaerli

## Abstract

Cell motility drives spatial pattern formation across diverse biological systems. Here, we engineer *Escherichia coli* motility in semi-solid agar to control Voronoi patterns in two and three dimensions, partitioning space into regions closest to their respective inoculation seeds. Consistent with our reaction–diffusion model, we observed that collisions between expansion fronts generate either biomass depletion (“gaps”) or accumulation (“anti-gaps”), governed by the relative diffusion rates of bacteria and nutrients. By engineering strains with distinct expansion rates and tuneable motility, and by integrating these experimental data into a dynamic Voronoi model, we achieved precise control over pattern geometry. This enabled the generation of gaps with varying widths, curved boundaries, asymmetric structures, seedless regions, and complex composite patterns. Together, these findings establish bacterial Voronoi patterns as a programmable platform for engineering multicellular spatial organization, with potential applications in synthetic biology and materials science.

## 1 Introduction

Fascinating polygonal patterns can be observed across multiple scales in biology, including giraffe’s skin patches [1], insect wing venation [2], honeycombs [3], plant cell walls [4], epithelial tissues [5] and trabecular bones [6]. Despite their structural and mechanistic diversity, they can be described using the same formalism: Voronoi diagrams. A Voronoi diagram is the partition of a plane or a volume into a series of regions, called Voronoi cells, based on the relative distance from a set of points, called seeds. Each Voronoi cell contains all the points that are closer to a given seed than to any other seed. The diagram appears as a mosaic of polygons, each of which represents the “area of influence” of a seed.

Despite informal use of Voronoi diagrams as early as the 17*^th^* century [7], they have became famous thanks to the work of Georgy Feodosievych Voronoy, who described the general n-dimensional case in 1908 [8]. Ever since, they have found countless applications across a variety of fields, ranging from meteorology [9] to medicine [10] and from robotics [11] to cell biology [12]. Voronoi-based structures are highly valued in tissue engineering as they display high resistance to mechanical stress [13, 14], low weight [15, 16] and efficient material usage [17]. They have therefore been used in biomedical lattices [14], cranial implants [15] and bone scaffolding [18]. However, the mechanisms underlying the development of Voronoi patterns in nature remain largely unknown [19].

Interestingly, bacterial communities can also generate Voronoi patterns, both in motile and non-motile contexts. For example, Voronoi diagrams have been used to study the role of competition during colonization [20] and the environmental factors influencing variation in colony size [21]. While the structure of surface colonies depends almost entirely on population growth, in soft agar, expanding colonies develop through the combined contributions of cell division and cell motility. If cells are inoculated at multiple sites, each inoculation point acts as a seed, and colonies expand until they reach the front originating from a neighbouring seed. This phenomenon partitions the agar into patches of high bacterial density, corresponding to Voronoi cells, separated by thin cells-depleted boundaries (gaps) [22–29]. Further lowering the agar concentration causes the two fronts to merge, a phenomenon that has been elegantly used to generate Voronoi tessellation with engineered *E. coli* expressing heterophilic cell-cell adhesins: when strains expressing complementary nanobody-antigen pairs merge, a straight line with high cell density forms at the interface [30].

Several hypotheses have been put forward to explain the gap between distinct colonies, with different mechanisms suggested for different species. In *Proteus mirabilis* swarms of two different strains, the gap is associated to self-nonself distinction and relies on short-range toxin-based killing mechanisms [24, 31]. Many bacteria, including *Bacillus subtilis*, encode similar mechanisms of kin discrimination [32, 33]. *E. coli* is also capable of inhibiting the growth of gram-positive and gram-negative bacteria, including other *E. coli* strains [28]. Theoretical work proposed that quorum sensing might play a role in the kin discrimination to avoid non-cooperative “cheaters” taking advantage of costly cooperative traits [34]. When the gap formation occurs between two identical strains, the repulsion was proposed to be caused by accumulation of toxic byproducts [21], or a change in pH [35, 36]. Work with *Pseudomonas putida* suggested that mechanical effects imposed by bacterial growth deformed the agar and formed a physical barrier [27]. The depletion of nutrients caused by growing colonies has also been used to justify the gap formation [25].

Here, we reveal that the ratio of bacterial to nutrient diffusion governs interface dynamics, driving the formation of either biomass “gaps” or “anti-gaps”. We demonstrate that bacterial expansion can be harnessed to engineer complex Voronoi tessellations in both two and three dimensions. By synergistically integrating modeling with genetic engineering, we achieve precise control over pattern geometry, enabling the generation of tunable gaps, curved boundaries, asymmetric structures, seedless regions, and complex composite patterns. Together, this work transitions the field from the observation of natural Voronoi patterns to the rational engineering of bacterial multicellular spatial organization.

## 2 Results

### Voronoi pattern formation and the role of effective cell diffusion

We first randomly seeded *Escherichia coli* MG1655 (WT) inside 0.35% LB agar. The resulting spatial organization consisted of polygonal sectors of varying size and shape, reflecting the initial distribution of inoculation points. As previously reported [22–29], we observed regions of high bacterial density corresponding to Voronoi cells, separated by thin, cell-depleted gaps (Figure 1a). Spatial image analysis confirmed that these domains closely followed a Voronoi tessellation (Figure 1a).

**Figure 1:**
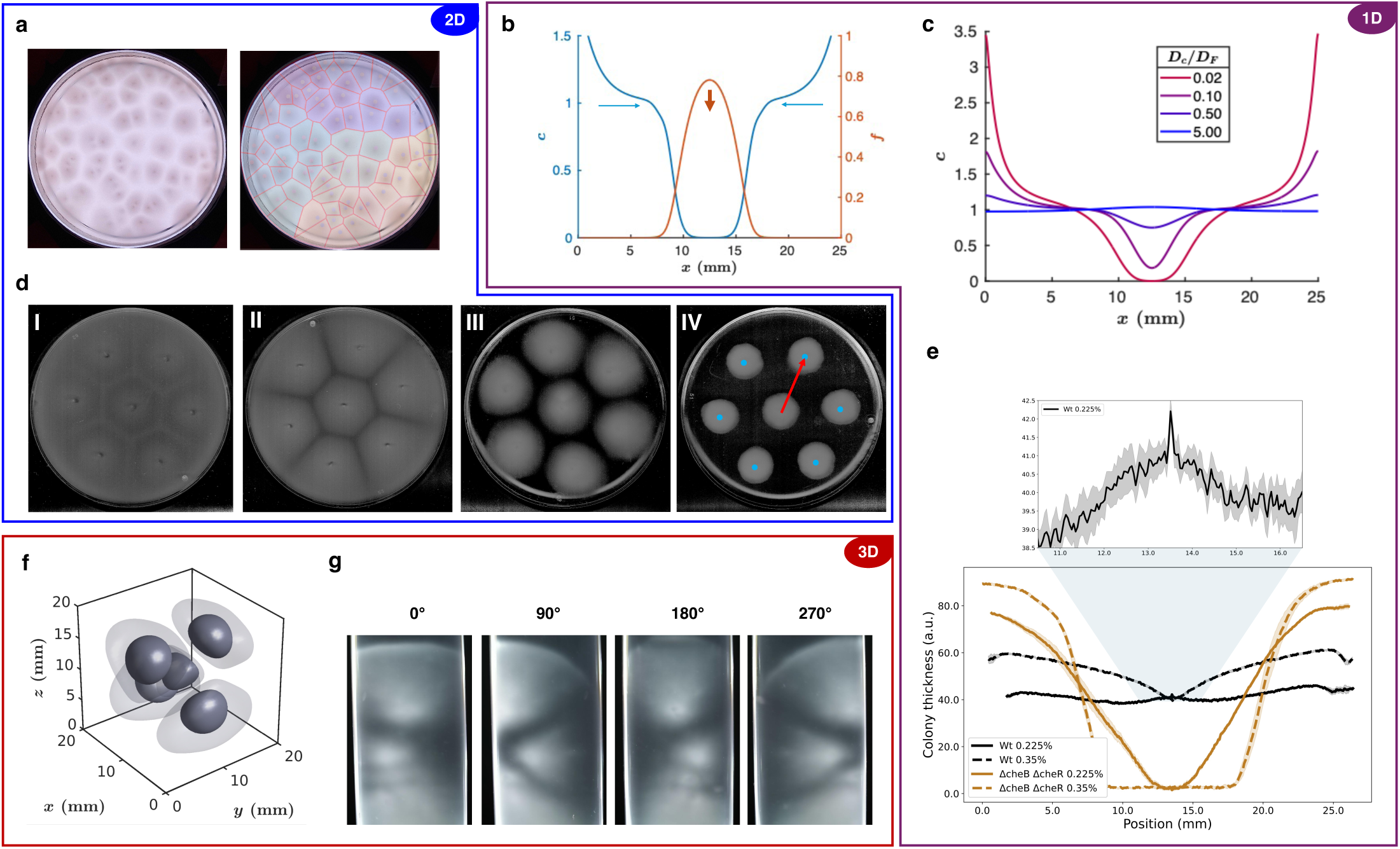
Motile bacteria form Voronoi patterns in low density agar. (a) *E. coli* MG1655 cells were randomly seeded in 0.35% LB agar and incubated for 24 h, leading to spatial partitioning between colonies (left). Processed image of the same plate highlights the Voronoi-like pattern (right). Gray dots indicate initial seed positions, and red lines show the corresponding Voronoi tessellation calculated from these positions. (b) One dimensional model illustrating nutrient consumption (orange, *f*) and bacterial expansion (blue, *c*) at *t* = 20 h. (c) Calculated gap profiles with the different effective diffusion coefficient ratios (*D_c_/D_f_*). (d) Hexagonal seeding of bacteria in LB agar imaged on a black background. Representative images of WT at 0.225% and 0.35% agar (I and II, respectively) and Δ*cheB* Δ*cheR* at 0.225% and 0.35% agar (III and IV, respectively). The red line indicates the path used to measure the gray-scale intensity profile from the center of the central colony to the center of one of the six neighboring colonies (blue dots). 6 measurements were performed for every plate. (e) Quantification of the gray-scale profiles shown in panel **d**. Lines represent the mean grayscale intensity of 6 measurements from the center colony to the neighboring colonies, with the standard error shown as shaded regions. The top graph shows a magnified view of the WT 0.225% agar condition (I), revealing a local increase in cell density between neighboring colonies, corresponding to the formation of the “anti-gap”. (f) Model of colonies expanding in three-dimensional space, demonstrating the emergence of Voronoi partitioning. (g) Experimental realization in a test tube, where immersed seeds expand and generate a 3D Voronoi pattern. Pictures were taken at different positions for better visualization, and the movie of the rotating tube is available as Supplementary Movie 3.

Subsequently, instead of seeding randomly, we inoculated the agar with bacteria by stabbing and pipetting 1 *µ*l of culture at seven specific locations (a central seed surrounded by six others arranged in a hexagonal pattern) and explored how changing agar concentration could affect the emergence of the gaps (Figure S1). At low agar concentrations (0.225%), we observed no gaps between the expanding colonies (Figure S1-I, Supplementary Movie 1). At intermediate concentrations (0.35%), regions of reduced cell density emerged between neighboring expanding colonies (Figure S1-II, Supplementary Movie 2), while at 0.45% agar, the effect was accentuated, enlarging the gap area (Figure S1-III). At 0.50% agar, bacterial motility was strongly impaired, likely due to reduced pore size, effectively suppressing cell motility (Figure S1-IV).

In these experiments, slower colony expansion consistently produced larger gaps, whereas faster expansion resulted in progressively smaller gaps, suggesting that the phenomenon is governed by a universal physical mechanism rather than a pathway-specific biological response. To identify this mechanism, we developed a phenomenological reaction–diffusion model (SI Section 4, *Mathematical modeling and numerical simulations*). The model describes bacterial dispersal as an effective active diffusion process that accounts for bacterial motility [37–39], cell proliferation through Monod kinetics, nutrient diffusion and consumption, and the gradual loss of motility as nutrients become depleted. The latter is implemented by making the effective cell diffusion coefficient dependent on the local nutrient concentration, allowing the colony to become stationary once nutrients are exhausted. It is important to emphasize that the cell diffusion coefficient should be interpreted as an *effective* transport coefficient rather than the microscopic random motility of individual bacteria. It summarizes the combined influence of multiple processes, including swimming speed, tumbling frequency, agar density, and other factors affecting colony expansion. Importantly, this effective diffusion coefficient can be estimated directly from experimentally measurable quantities, namely the colony expansion velocity and growth rate.

In the dimensionless formulation of the model, nutrients are converted into biomass, such that the sum of the cell and nutrient concentrations represents the total amount of material available at a given position. As shown in SI Section 3, the travelling-wave solution naturally develops either a positive or a negative deviation of this sum from its bulk value (Figure S3). This effect originates from the sigmoidal shapes of the cell and nutrient profiles: one profile increases while the other decreases across the reaction front, and their shapes and relative positions determine whether their sum exhibits a local minimum or maximum (Figure 1b). When two travelling fronts collide, this transient surplus is preserved and gives rise to either a gap or an accumulation of bacteria at the boundary between neighboring colonies (Figure 1c). The analytical treatment further shows that, the sign of this surplus is determined solely by the relative magnitudes of the effective cell and nutrient diffusion coefficients. When the effective cell diffusion coefficient is smaller than the nutrient diffusion coefficient, the travelling front develops a local depletion, producing a gap after collision. Conversely, when the effective cell diffusion exceeds nutrient diffusion, the front contains a local surplus that generates an accumulation of bacteria. The magnitude of the surplus is predicted to increase with the difference between the two diffusion coefficients.

A key prediction of the model is that under certain conditions instead of gaps, we should be able to observe an accumulation of bacteria (“anti-gaps”) at the contact zone. The theoretical results suggest that gap and “anti-gap” formation is a generic consequence of the structure of the travelling reaction front. At the phenomenological level, the effect is governed by the effective transport of cells relative to nutrients. At the microscopic level, however, this effective transport reflects the combined influence of bacterial motility, agar density, nutrient availability, and other biological and physical factors that regulate colony expansion.

To test these predictions, we decided to analyse the experimental patterns in plates photographed against a black background, which improved the visualization of local variations in cell density: the observed brightness relative to the dark background arises from light scattering by the bacteria and therefore provides a proxy for local cell density. Importantly, while at 0.35% (Figure 1d-II) we observed again gaps between neighbouring Voronoi regions, at 0.225% agar we could detect small local maxima of bacterial density (Figure 1d-I), as quantified by the measurement of the gray-scale profile obtained from the images of the expanding colonies (Figure 1e). Such a transition from a depletion to accumulation of bacteria is difficult to reconcile with a mechanism based solely on the accumulation of inhibitory compounds or the depletion of nutrients, but it is consistent with our theoretical predictions. Indeed, the experimentally determined effective diffusion coefficients presented in SI Section 4 are in agreement with an effective cell diffusion coefficient that exceeds the nutrient diffusion coefficient in the agar (Figure 1e).

We also tested the *E. coli* MG1655 ΔcheB ΔcheR strain, which has been reported to exhibit significantly impaired motility [40], to investigate how reduced bacterial motility affects colony expansion across different agar concentrations (Figure 1d-III-IV,e). As expected, the mutant strain had much lower motility rate and consequently could not colonize the entire plate even at 0.225% agar concentration. Notably, the model predicted that this low-motility phenotype also enables the formation of gap Voronoi patterns in three dimensions (Figure 1f). To experimentally confirm this prediction, we embedded slow-migrating *E. coli* MG1655 ΔcheR ΔcheB in 0.30% LB agar within a test tube. Under these conditions, colony expansion became sufficiently slow to observe three-dimensional high-density regions surrounded by low-density regions (Figure 1g).

### Weighted Voronoi patterns driven by front speed variation

Voronoi domains formed by colonies with similar expansion rates exhibit straight boundaries at their interfaces. However it has been observed that different species with distinct expansion rates can lead to asymmetric competition, whereby faster-expanding colonies displace slower ones, resulting in curved interfaces characteristic of weighted Voronoi tessellations [29]. To experimentally control this effect within the same species, we first characterised three *E. coli* strains with distinct metabolic burdens, which ultimately affects their colony expansion rate: *E. coli* MG1655 wild-type (WT) displayed the highest expansion rate of the three strains. The second strain (TS-purple) carried a plasmid constitutively expressing the purple chromoprotein tsPurple [41] and exhibited a slightly slower expansion, but very similar to that of the WT. The third strain (Amil-CP) was transformed with a plasmid for constitutive expression of the blue chromoprotein amilCP [41] and showed the slowest expansion, consistent with an increased metabolic burden (Figure 2a). We attribute the reduced expansion of the Amil-CP strain to the non–codon-optimized amilCP gene (from *Acropora millepora* [42]).

**Figure 2:**
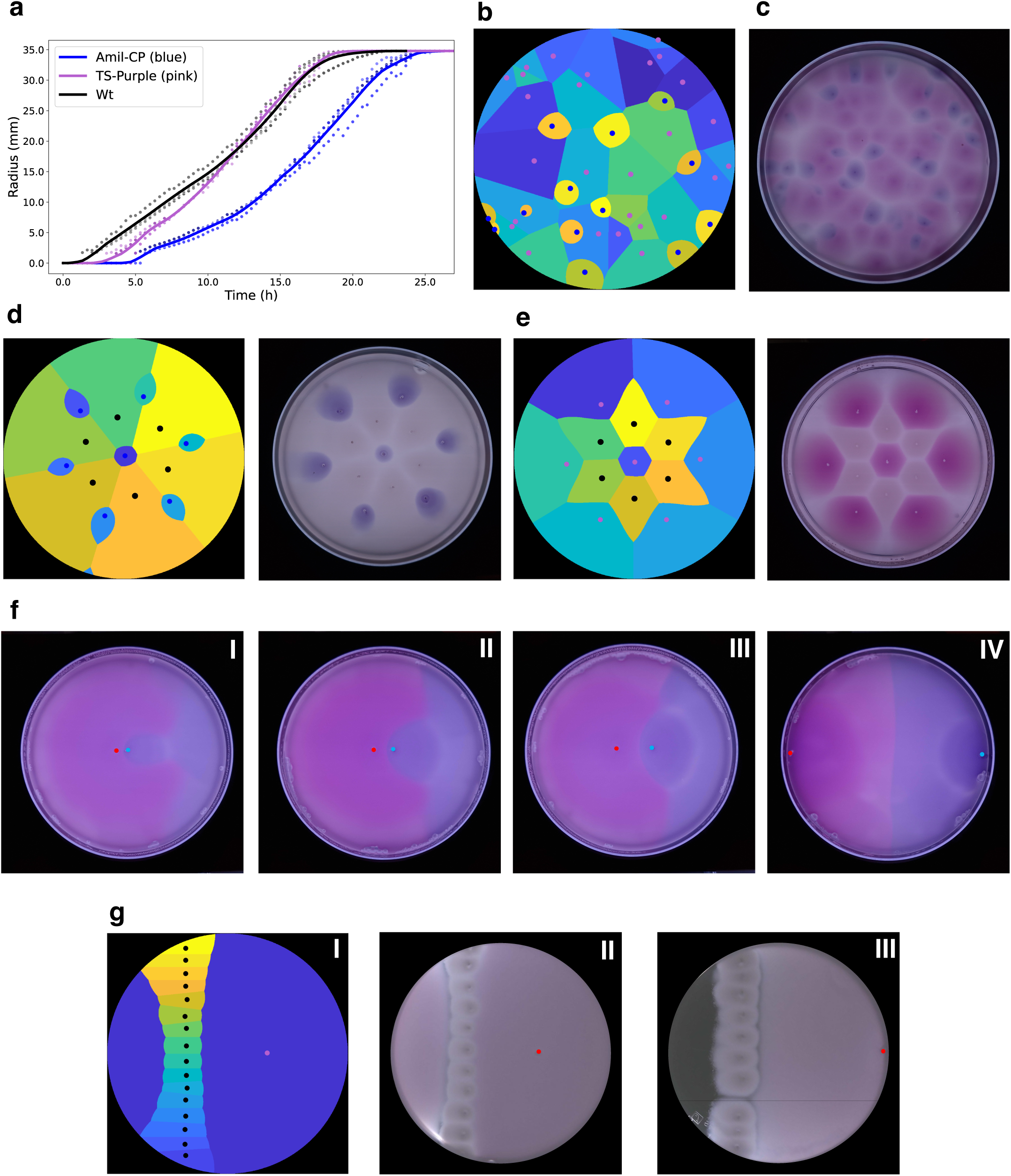
The relative expansion rates of neighboring colonies determine the shape of their boundary. (a) Expansion dynamics of the *E. coli* MG1655 strain (WT - black), and the same strain harboring Amil-CP (blue) or TS-Purple (pink) plasmids. Dots represent biological replicates (n=3), solid lines represent smoothed averages. (b) Dynamic Voronoi model of a weighted Voronoi pattern, produced by random seeding. Colonies with different expansion rates generates curved boundaries: faster-expanding seeds (pink seeds) dominate slower ones (blue seeds). (c) Random seeding of Amil-CP (blue) and TS-Purple (pink) strains on LB 0.35% agar. The faster-expanding TS-Purple colonies deform the boundary, curving into the slower Amil-CP regions. (d-e) Seeds inoculated at identical positions on LB 0.35% agar, with the dynamic Voronoi model (left image) and experiment (right). Pattern morphology depends on relative expansion rates: (d) a weighted Voronoi pattern emerges between WT (white) and Amil-CP (blue), (e) a classical Voronoi pattern is observed when expansion rates are similar (WT, white; TS-Purple, pink). (f) TS-Purple and Amil-CP strains inoculated (red and blue dots, respectively) at increasing distances from one another (I to IV). At close distances, the TS-Purple’s expansion rate is higher than Amil-CP’s, resulting in a weighted Voronoi pattern. At larger distances, expansion rates of both strains are similar, leading to straight boundaries. (g) I: Dynamic Voronoi model illustrating the emergence of a seedless region in a weighted Voronoi configuration. II: experimental demonstration of seedless region formation depending on the spacing between the slowly expanding ΔcheBΔcheR strain (barrier) and the fast-expanding TS-Purple seeding. III: Increasing the distance between the barrier and the seed leads to the formation of a exclusion zone (LB 0.225% agar).

While our reaction–diffusion model offers mechanistic explanatory power, its complexity limits its applicability for rapid quantitative pattern design when only simple experimentally measurable quantities are available. To overcome this limitation, we developed a second approach, what we call the dynamic Voronoi model, which predicts the final colony partition directly from experimentally measured expansion curves (SI Section 4, Mathematical modeling and numerical simulations, II). In this approximation, each inoculation site is treated as an expanding source whose front propagation is determined by its individual radial growth law, and each spatial position is assigned to the colony that reaches it first. The expansion rate of each inoculation site determines the weight that influences the distance computation. This generates a generalized weighted Voronoi tessellation in which boundaries are determined by front arrival times rather than purely geometric distances. Because this minimal method requires only simple colony expansion measurements and avoids solving the full reaction-diffusion dynamics, it provides a computationally efficient and quantitatively robust framework for forward design of target spatial arrangements.

Using this dynamic Voronoi model, we predicted weighted Voronoi patterns (Figure 2b, Figure S2a) between the Amil-CP and TS-Purple strains. To test this predictions, we first seeded these strains randomly. Indeed, the colonies of the blue coloured strain (slower) were bent by the fast expanding purple strain (Figure 2c). Consequently, we were able to predict and produce distinct patterns, depending on the combination of strains (and expansion rates) used, even when the initial seeding arrangement was identical (Figure2d,e, Figure S2b,c).

It is worth noticing that the expansion rate differences were transient. After 15 h, the expansion rate of the Amil-CP strain approached that of WT and TS-purple (Figure 2a). As a result, the geometry of the resulting patterns depends on the distance between inoculation sites: at short inter-seed distances, early-time expansion asymmetries dominate, yielding weighted Voronoi geometries with curved boundaries, whereas at larger separations, later-time dynamics prevail, resulting in classical Voronoi patterns with straight interfaces (Figure 2f, Figure S2e). Thus, seeding distance between colonies with different expansion rates can be combined to produce unconventional patterning that changes the boundary shape as the pattern evolves (Figure S2f, Figure S2e).

The combination of expansion rates and seed positions can also modulate patterning through a second mechanism intrinsic to weighted Voronoi systems: the emergence of regions devoid of any seed. In our model, this arises when a slower-expanding colony forms a barrier that interrupts the continuity of a faster one (Figure 2g-I, Figure S2d). Experimentally, we reproduced this behaviour using the slow-migrating MG1655 ΔcheR ΔcheB strain and the fast Ts-purple strain. The slow strain allowed the faster strain to pass through, creating a region devoid of a seed, yet occupied by the TS-purple strain (Figure 2g-II).

Interestingly, increasing the initial distance between the fast and slow strain seeds experimentally revealed a qualitatively distinct behaviour: When the slowly expanding, high-density colonies encounter one another before the faster-expanding strain arrives, they effectively form a barrier that blocks subsequent penetration. As a consequence, an isolated uncolonised region can persist on the opposite side, giving rise to a region devoid of bacteria (Figure 2g-III).

It is important to note that the reaction–diffusion model does not permit the formation of such permanently excluded regions, provided that nutrient availability is sufficient to support growth across the entire plate. Under these conditions, the model predicts eventual occupation of all accessible space (Figure S2d). An analogous limitation applies to the dynamic Voronoi model: by definition, every point in the domain is assigned to at least one Voronoi cell, such that true empty regions cannot emerge, although individual Voronoi territories may become spatially discontinuous (Figure 2g-I,II).

Overall, we demonstrated that the interplay between expansion rate and seed positioning can be tuned to generate a wide range of spatial patterns.

### Patterning control with CheZ levels

We demonstrated that metabolic burden influences colony expansion (Figure 2), which in turn affects the Voronoi pattern. To further extend control over pattern formation, we set out to regulate motility directly. Among the various regulators involved in the control of the motility, the *che* genes play a key role: the protein CheY in its phosphorylated form binds to the flagellar motor and induces a change in rotation direction, from counterclockwise (run) to clockwise (tumble). The balance between the activities of the phosphatase CheZ and the kinase CheA determines the amount of phosphorylated CheY, ultimately resulting in *E. coli* ability to swim randomly or directionally [43]. Motility of bacteria in soft agar requires both running and tumbling, suggesting that tumbling is necessary for cells to ‘back away’ from obstructions in the agar [22, 44]. Consequently, motility in soft agar is a non-monotonic function of the ratio of CheA to CheZ. Overexpression or underexpression of CheZ disrupts the optimal signaling sensitivity, leading to reduced motility in soft agar, creating a bell-shaped response [45, 46].

To systematically control the diversity of patterns arising from differences in expansion rate, we thus sought to tune motility through inducible expression of CheZ. We generated an *E. coli* MG1655 ΔcheZ strain complemented with plasmids in which CheZ expression is driven either by the arabinose-inducible pBAD promoter or by the quorum-sensing–responsive pLux promoter (Figure 3a, d). Induction with L-arabinose increased motility in a dose-dependent manner, with time-lapse imaging confirming that expansion rates can be precisely tuned as a function of inducer concentration (Figure 3b). Localized addition of L-arabinose (2 *µ*L) generated a diffusive gradient, resulting in spatially heterogeneous expansion: colonies proximal to the inducer occupied the entire Voronoi cell, whereas distal colonies remained almost non-motile and only occupied a small portion of the Voronoi cell (Figure 3c).

**Figure 3:**
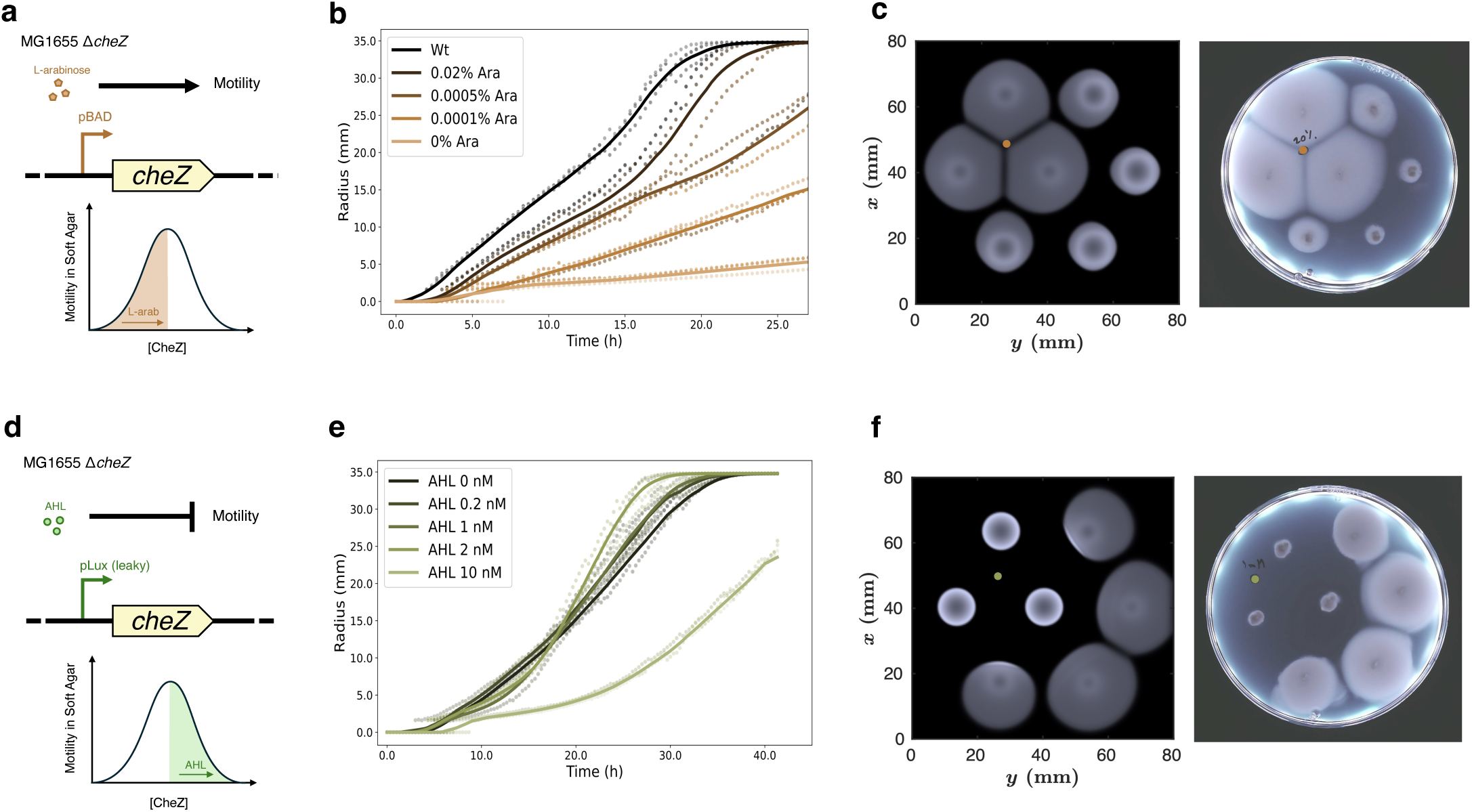
CheZ expression modulates bacterial motility, enabling the generation of asymmetric Voronoi patterns. (a) Schematic representation of the L-arabinose-inducible cheZ circuit. The level of motility is directly proportional to the inducer concentration. (b) Expansion rates of WT and pBAD-cheZ strains at different arabinose concentrations. Dots represent biological replicates (n=3), solid lines represent smoothed averages, LB 0.35% agar. (c) Left: simulation (*t* = 15 h) based on fitted parameters collected from ‘b’. Right: A 2 *µ*L drop of 20% L-arabinose was added at the delimited spot. As the inducer diffuses across the plate, seeds show increased motility the closer they are to the inducer drop, LB 0.35% agar. (d) Schematic representation of the AHL-inducible cheZ circuit. The level of motility is inversely proportional to the inducer concentration. (e) Expansion rate of AHL-inducible-cheZ strain at different AHL concentrations. Dots represent biological replicates (n=3), solid lines represent smoothed averages., LB 0.35% agar. (f) Left: simulation (*t* = 15 h) based on fitted parameters collected from ‘e’. Right: A 2 *µ*L drop of 1 mM AHL was added at the delimited spot. As the inducer diffuses across the plate, seeds show decreasing motility the closer they are to the inducer drop, LB 0.35% agar.

On the other hand, the pLux-cheZ construct, characterized by basal leakiness, conferred near–wild-type motility in the absence of induction (Figure 3d-e). Upon addition of the inducer N-acylhomoserine lactone (AHL, 3-oxohexanoyl-homoserine lactone), *cheZ* overexpression reduced motility, reversing the spatial response: colonies near the inducer showed limited expansion, while distant colonies expanded more readily and occupied the entire Voronoi cell (Figure 3f).

### Gap filling in composite Voronoi patterns

The self-organized formation of Voronoi tesselations presents a new approach to spatial patterning in engineered living materials. For some applications, it may be advantageous to assign a distinct function to the regions between Voronoi sectors rather than leaving these gaps unoccupied. This led us to exploit the “anti-gap” phenomenon observed at low agar concentrations (Figure 1e-IV). If a faster expanding population can preferentially colonize the gap regions before the neighboring colonies meet, it may be possible to generate composite patterns in which different bacterial populations occupy distinct spatial domains within the same Voronoi architecture.

Building on this premise, we explored co-inoculating a fast-growing and a slow-growing strain at the same site. We inoculated a mixture of TS-Purple and Amil-CP strains in LB 0.225% agar. While the slow-growing Amil-CP strain remained confined to the Voronoi domains, the fast growing TS-purple strain rapidly colonised the gap regions, producing a mesh-like pattern (Figure 4a-I-II).

**Figure 4:**
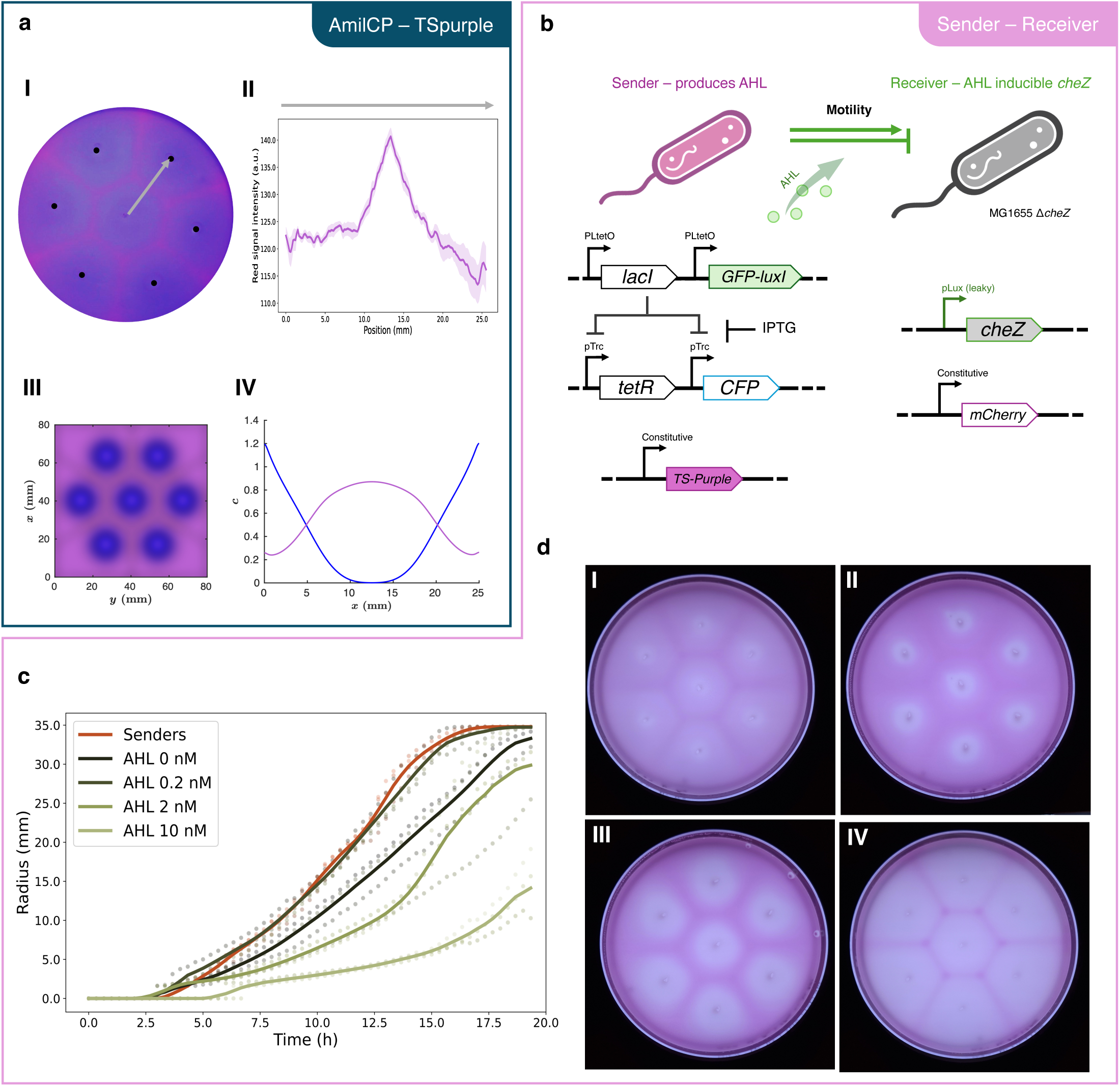
Co-inoculation of fast- and slow-expanding strains generates composite Voronoi patterns, with each strain predominantly occupying distinct regions. (a) I: Composite Voronoi pattern where the seed areas are largely dominated by Amil-CP strain, while the gap regions are colonised by TS-Purple strain; LB 0.225% agar. The gray arrow indicates the path used to measure the red color intensity, from the center of the central colony to the center of one of the six neighboring colonies (black dots). II: Red color intensity from the composite Voronoi pattern in ‘I’. Solid red line represents the average of six measurements from the center colony to each neighbor colony, and shaded area indicates standard error of the mean. III: 1D model depicting the density of two populations, faster-expanding in pink and slower-expanding in blue. IV: 2D reaction–diffusion model of the composite Voronoi. (b) Schematic showing the toggle switch circuit in the Sender strain and the AHL-controlled motility (via cheZ) in the Receiver strain. (c) Expansion rate of the Sender strain and the Receiver strain at different inducer concentrations. Dots represent biological replicates (n=3), solid lines represent smoothed averages, LB 0.225% agar. (d) The width of the gaps occupied by Sender cells can be tuned by varying IPTG and AHL concentrations in LB 0.20% agar: I - No inducers, II - IPTG 1mM + AHL 2nM, III - IPTG 1mM + AHL 1nM, IV - IPTG 1mM.

Our reaction–diffusion simulations reproduce this spatial segregation, with the faster-dispersing strain dominating the expanding front and the slower-dispersing strain becoming enriched in the interior (Figure 4a-III, IV). The model indicates that the spatial distribution of the two strains is determined by two competing effects: During the early stages of colony growth, nutrients are abundant, and both strains proliferate close to their maximal growth rates. Under these conditions, the larger effective diffusion coefficient of the TS-Purple strain results in a faster expanding front, allowing it to colonize the outer regions of the colony and the boundaries between neighboring Voronoi domains. Behind the advancing front, however, nutrient concentrations gradually decrease. Because the Amil-CP strain has a smaller Monod half-saturation constant (SI Section 4.3, Figure S5), it is able to sustain growth at lower nutrient concentrations and gradually outcompete the TS-Purple strain in the interior of the Voronoi cell.

To tune the size of the regions occupied by the different strains in these composite patterns, we switched to different strains: a Sender strain that modulates the motility of a Receiver strain in a controllable manner. Specifically, the Sender strain harbored a toggle-switch circuit regulating AHL production [47], while the Receiver strain contained the pLux-cheZ circuit and a constitutive mCherry reporter (Figure 4b). In the bistable toggle-switch, the expression of *lacI* and *gfp-luxI* (an AHL synthase fused to the GFP reporter) is repressed by TetR, while *tetR* expression is itself repressed by LacI. In the absence of IPGT (isopropyl *β*-d-1-thiogalactopyranoside), the circuit preferentially assumes the LacI/GFP-LuxI state, leading to AHL synthesis. Upon addition of IPTG, the switch transitions to the TetR state, resulting in repression of *luxI* and cessation of AHL production (Figure 4b). Additionally, we labelled the Sender strain with the TS-Purple plasmid (appearing pink). We did not utilize the fluorescent reporters (GFP and mCherry) in either strains for imaging.

We first characterized the expansion rates of the Sender and the Receiver strains at different AHL concentrations in LB 0.225% agar (Figure 4c). The Sender strain expanded faster than the Receiver strain in the absence of AHL, while the Receiver exhibited a more graded response to AHL induction at this lower agar concentration than in 0.35% (Figure 3e). Interestingly, at 0.2 nM AHL, the Receiver displayed increased motility relative to the uninduced state, effectively spanning both sides of the bell-shaped response curve (Figure 4c).

We then mixed and inoculated the Sender and Receiver strains together at fixed positions to investigate whether their spatial organization could be controlled through the signaling circuit. In the absence of inducer, no clear composite pattern emerged, apart from the previously observed accumulation at the contact zone between individual Voronoi regions, which was particularly visible due to the pink pigmentation of the Sender strain (Figure 4d-I). In contrast, addition of 1 mM IPTG produced a composite pattern, in which the pink Sender cells were predominantly confined to the gap regions, while the white Receiver cells preferentially colonized the Voronoi sectors (Figure 4d-IV). We then additionally supplemented the 1 mM IPTG with 1 or 2 nM of AHL (homogeneously distributed in the plate) to further decrease expansion rates of the Receiver strain. This increased the width of the gap regions occupied by the Sender bacteria, demonstrating that the spatial organization of the two populations can be controlled through the interaction of their motility rates (Figure 4d-II-III).

Together, these results indicate that differential motility and the resulting density asymmetry are key determinants of this composite pattern, enabling stable partitioning in which one strain occupies the Voronoi cells and the other fills the gaps.

## 3 Discussion

In this study, we leveraged the capacity of *E. coli* to move inside soft agar to generate Voronoi patterns. The approach leads to the tessellation of the agar into polygonal shapes, each originating from an inoculation seed, in both two and three dimensions (Figure 1). Bacterial Voronoi patterns in agar plates have attracted considerable interest due to the characteristic gaps that emerge between neighboring colonies [20, 21, 29, 30]. Several mechanisms, including byproduct accumulation, chemotaxis, and nutrient depletion, have been proposed to explain Voronoi patterns across bacterial species and experimental conditions [21, 24, 25, 28, 31–33, 35, 36], however, a universal framework remained elusive.

Building on previous studies on bacterial motility and Voronoi pattern formation [9, 25, 48], we established a phenomenological reaction–diffusion model describing bacterial motility, growth, and nutrient consumption. The model predicted that the propagating reaction front develops either local biomass depletion or accumulation, depending on the relative effective diffusion of cells and nutrients. When neighboring colonies collide, this imbalance is preserved at the collision boundary, producing either a cell-depleted gap or, when cell diffusion exceeds nutrient diffusion, a cell-enriched “anti-gap”. Indeed, consistent with our theoretical predictions, we detected small local maxima in bacterial density at the contact zones of Voronoi patterns at 0.225% agar (Figure 1d–e). To the best of our knowledge, this naturally occuring accumulation has not been previously reported in bacterial Voronoi patterns. Such a transition from bacterial depletion to accumulation is difficult to reconcile with mechanisms based solely on the accumulation of inhibitory compounds or nutrient depletion. However, our model indicates that this phenomenon arises from the coupled transport of cells and nutrients within the traveling reaction front.

Whereas previous studies have focused on observing Voronoi patterns that arise naturally following inoculation, our work actively controls these patterns. By genetically tuning bacterial motility, we generated a diverse range of Voronoi patterns, including curved boundaries and asymmetric organization, colonization of initially unseeded regions, barriers, and composite patterns in which specific strains are localized to the gap regions. These designs were supported by the dynamic Voronoi model that allowed us to quickly and accurately predict Voronoi patterns based on the input positions and expansion rates of the involved strains.

Specifically, by seeding strains with different expansion rates (Figure 2), we generated curved boundaries rather than straight boundaries, successfully realizing weighted Voronoi patterns, including the emergence of seedless regions and barriers (Figure 2g). In addition, we directly controlled motility by tuning the expression of CheZ to create asymmetric Voronoi diagrams (Figure 3). Furthermore, co-inoculation of two strains at a single site yielded previously unreported novel patterns characterized by ‘gaps’ occupied by the first strain and sectors filled by the second, slow-growing strain (Figure 4a). Finally, we demonstrated the ability to dynamically tune the relative areas occupied by the two strains, leveraging the interaction between a Sender strain, modulating the motility of a Receiver and inducer concentration. Our work complements existing efforts to engineer (self-organized) spatial patterns by controlling bacterial motility [26, 30, 49–52].

Recent work suggested that Voronoi patterns embody a universal geometric principle, providing a robust and reproducible framework for organization both in microbial communities and multicellular organisms [29]. The significance of the spatial organization we have studied here within semi-solid matrices is particularly evident in microbial communities found in mucosal environments, such as the gut, where microorganisms compete for space and resources. Motility helps bacteria move to new surfaces, after which they can form stationary biofilms. Our findings suggest that differences in bacterial expansion dynamics can contribute to the emergence of complex spatial organization. For example, even less motile populations can form physical barriers that restrict neighboring colonies from accessing new territories, providing a mechanism for spatial exclusion without direct antagonism. Our composite Voronoi patterns demonstrate how multiple populations with different expansion rates can self-organize to occupy distinct regions. It is intriguing to speculate that such spatial organization could provide functional advantages in confined environments, including efficient biomass distribution and enhanced resistance to mechanical stress [13, 14, 53].

Beyond their relevance to natural microbial communities, we believe that microbial Voronoi patterns hold significant potential for engineering living materials [54]. For instance, the barrier formation (Figure 2g-III) may provide useful for applications that require empty channels or confined growth compartments. Co-inoculation of a fast-expanding, biopolymer-producing strain with a slower, non-polymer-producing strain could generate composite Voronoi patterns in which the polymer-producing bacteria forms a structural framework confined to the gap regions. Similarly, functionalizing the strain present in the gap areas with gold nanoparticles could confer electrical conductivity [55], transforming the resulting pattern into a network of biological “wires” and providing the basis for circuit-like living materials with programmable electrical properties. Alternatively, by promoting biomineralization of 3D Voronoi patterns one could create three-dimensional tunable porous structures with high strength and toughness, mimicking natural materials such as nacre [56] and bones [18]. We believe that our simple yet innovative Voronoi pattern platform could inspire new approaches and accelerate the development of the next generation of bio-based materials.

## 4 Methods

### Strains and plasmids

All the experiments performed in this study used strains derived from *E. coli* str. K-12 substr. MG1655 (NCBI accession number *NC* 913000). All strains used in this work are listed in Table 1 in SI Section 5. *E. coli* motility mutants (MG1655 ΔCheZ and MG1655 ΔCheBΔCheR) were constructed using the *λ* Red recombination protocol [57]. Briefly, primers carrying overhangs homologous to the regions upstream and downstream the target genes were used to amplify a kanamycin resistance cassette flanked with FRT sites from the template plasmid pKD13. Co-transformation of the linear construct with pKD46, a plasmid expressing the *λ* components (*γ*, *β*, *exo*), resulted in homologous recombination between the target gene and the antibiotic resistance, allowing for selection of mutants. The kanamycin cassette was subsequently removed via expression of the Flp flippase from the helper plasmid pCP20, and strains were checked for loss of antibiotic resistance. Mutants were checked via colony PCR using primers binding to the regions upstream and downstream the target genes. As *cheB* and *cheR* are adjacent on *E. coli* genome, we deleted them simultaneously with one single construct.

DNA fragments were amplified via PCR using Phanta Max Master Mix (Vazyme) for cloning and Taq Max Master Mix (Vazyme) for sequencing. DNA fragments were assembled using NEBuilder HiFi DNA Assembly Master Mix (NEB). All plasmids and primers used in this work are listed in Tables 2 and 3 in SI Section 5, respectively. For the L-arabinose-inducible cheZ, the genomic sequence of *cheZ* was first cloned into a pCDF vector. Subsequently, a copy of the *araC* regulator and the pBAD promoter were placed upstream of *cheZ*. For the AHL-inducible cheZ, we received two plasmids, pLux-B0031-cheZ-(J23100+B0030+mCherry+B1002) and pLuxR(35k)-LasI(20K), kind gifts from Jiandong Huang [51]. pLux-B0031-cheZ-(J23100+B0030+mCherry+B1002) is a p15A ori vector that carries a copy of the *cheZ* gene under control of a pLux promoter and constitutively expresses mCherry from a pJ23100 promoter. We included at the beginning of the *cheZ* sequence an additional ATG codon, to make it identical to the genomic sequence, generating the pLux-cheZ-mCherry plasmid. Plasmid pLuxR(35k)-LasI(20K) is a pMB1 ori vector constitutively expressing the regulator *luxR*, alongside the quorum sensing synthase *lasI* (irrelevant for this work). The pLux-cheZ-mCherry and the pLuxR-LasI plasmids were co-transformed into the same strain, in order to tune *cheZ* expression via addition of AHL. The Toggle Switch - LuxI plasmid for inducible production of AHL was constructed previously in our group (Addgene ID #251147) [47]. Chromoprotein expressing plasmids, amilCP (Addgene ID #117847) and tsPurple (Addgene ID #117848), were a kind gift from Anthony Forster [41].

### Agar plate assays

Bacteria were grown overnight in LB liquid medium at 37 °C. Next, 1 *µ*l of overnight culture was stabbed into semi-solid agar plates (25 mL in a 90mm x 15mm Petri dishes). For random bacterial seeding, overnight cultures were serially diluted to an estimated concentration of approximately 5 bacteria/*µ*L, and 20 *µ*L of the diluted culture was added to molten LB agar maintained at approximately 37 °C. Agar plates were freshly made on the same day of the experiment, with densities from 0.20% to 0.50%, and antibiotics when necessary (chloramphenicol 30 *µ*g/ml, kanamycin 50 *µ*g/ml, spectinomycin 50 *µ*g/ml). Plates were incubated upright at 37 °C for 24 to 48 h.

### Agar plates image acquisition

To collect timelapses of expanding bacterial colonies, we used an office scanner (Epson Perfection V39II) to scan the plates every 20 minutes inside the incubator. An automatic clicker code was used to trigger the scans. The lids of the Petri dishes were painted with black primer (Warpaint) to create a black background and decrease reflection.

For timelapses and single pictures showing more detailed images of the wave expansions or the final images for comparison, we used the ‘Rocket’, a custom-made imaging platform consisting of a high light sensitivity commercial camera (ELP-USB130W01MT-BFV) and a Petri dish holder surrounded by a white LED ring, all coated with a black 3D printed shell to shield from external light sources. The plate was incubated in the Rocket facing up and was imaged from above.

### 3D Voronoi pattern

An overnight culture of the MG1655 ΔcheR ΔcheB strain was diluted 10^6^-fold and 1 *µ*L was inoculated in the test tube containing 3 mL of non-solidified LB 0.30% agar and after solidification, it was incubated at 37 °C for 48 h. After the incubation, a stepper motor controlled via Arduino Uno microcontroller was used to turn the test tube and take pictures with a commercial camera (ELP-USB130W01MT-BFV).

### Composite Voronoi experiment

For the composite Voronoi assay, strains were grown overnight in liquid LB medium. 1 *µ*L of an equal mixture of both strains (TS-Purple and Amil-CP, or Sender and Receiver) was stabbed into freshly prepared LB 0.225% agar (with the necessary antibiotics), at the appropriate locations. For the Sender-Receiver experiments, IPTG and/or AHL were added during the preparation of the LB 0.20% agar plates. Plates were incubated facing up at 37 °C for 24 h, separated by empty plates.

### Mathematical modeling and numerical simulations

#### I Reaction–diffusion model

To describe and understand the mechanisms underlying the experimental observations, we developed a phenomenological reaction– diffusion model. The model captures the coupled dynamics of bacterial growth, nutrient consumption, and cell dispersal, while remaining sufficiently simple to identify the physical origin of gap and antigap formation. Bacterial proliferation and colony expansion is described by a Fisher-type reaction–diffusion equation with Monod kinetics, whereas nutrient consumption is directly coupled to biomass production through the same growth term. Since the bacterial strains are motile, cell dispersal is described using an effective diffusion model developed for active matter [37–39], in which the diffusion coefficient depends on the local nutrient concentration and, in some cases, on small diffusible signaling molecules such as AHL or L-arabinose. As nutrients are depleted, the active motility gradually ceases, leading to an immobilized colony after nutrient exhaustion. In SI Section 2 we introduce dimensionless concentration variables, which eliminate the nutrient-to-biomass conversion factor and yield the following governing equations:

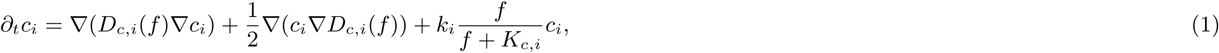

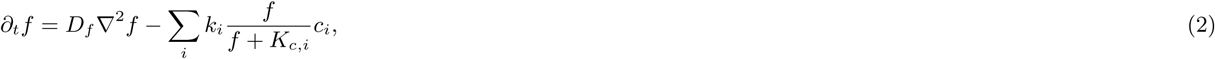

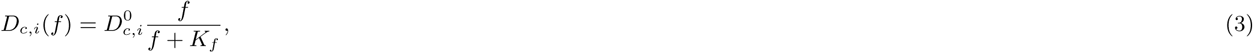

where 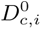 is the maximum effective diffusion coefficient of cell type *i*, *k_i_* is its proliferation rate, and *K_c,i_* is the Monod half-saturation constant for growth. The parameter *D_f_* denotes the nutrient diffusion coefficient, while *K_f_* determines the nutrient concentration at which the effective cell diffusion is reduced to half of its maximum value. Unless stated otherwise, we used *D_f_* = 0.85 mm^2^*/*h for small molecules or peptides in low-density agar [58] and *K_f_* = 0.01.

The system of reaction–diffusion equations was solved numerically using finite-difference methods in one-, two-, and three-dimensional domains. One-dimensional simulations were performed both in Cartesian and polar coordinates, while two- and three-dimensional simulations used Cartesian grids. Spatial derivatives were approximated with second-order central finite differences, and time integration was performed using an explicit Euler scheme. Zero-flux (Neumann) boundary conditions were imposed for all variables, corresponding to an isolated system without exchange across the domain boundaries.

Unless stated otherwise, simulations were performed on domains of length *L* = 25 mm, 80 mm, and 20 mm in one, two, and three dimensions, respectively. The spatial resolution was *h* = 0.025 mm in one dimension and *h* = 0.4 mm in two and three dimensions. The maximum simulated time was *T* = 200 h in one dimension, *T* = 30 h in two dimensions, and *T* = 200 h in three dimensions, with time steps of Δ*t* = 4 *×* 10*^−^*^5^ h, 0.003 h, and 0.02 h, respectively. Simulations were terminated earlier if the nutrients were completely depleted and the system had reached a stationary state. The nutrient concentration was initialized uniformly throughout the domain according to *f* (**x**, 0) = 1. Bacterial colonies were initialized as localized super-Gaussian distributions,

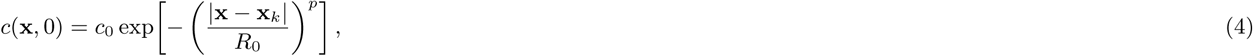

where *c*_0_ denotes the initial colony density, *R*_0_ the characteristic colony radius, and *p* controls the sharpness of the profile. We used *c*_0_ = 1 in one and three dimensions and *c*_0_ = 0.1 in two dimensions. The initial colony radius was *R*_0_ = 3 mm in two dimensions and *R*_0_ = 2 mm in three dimensions. The vectors **x***_k_* specify the colony centres, which were positioned according to the corresponding experimental configuration.

For simulations including externally supplied signaling molecules, such as arabinose or AHL, we used *p* = 2, corresponding to a Gaussian initial concentration profile representing diffusion from a localized source. The effective cell diffusion coefficients in the presence of L-arabinose and AHL were determined from measurements of colony expansion at different signaling molecule concentrations (Figure 4c). The corresponding diffusion coefficients were then interpolated to obtain the values used for the concentrations occurring in each simulation. L-arabinose and AHL initial concentration profiles had a characteristic radius of *R*_0_ = 15 mm, with peak concentrations of *c*_0_ = 7 *×* 10*^−^*^5^, % for arabinose and *c*_0_ = 100 nM for AHL. The cell-specific model parameters and the corresponding experimental measurements used for their estimation are summarized in SI Section 4.

#### II Dynamic Voronoi model

As a reduced predictive approximation, we additionally constructed spatial competition patterns directly from experimentally measured colony expansion curves using a generalized weighted Voronoi algorithm. We refer to this approach as “Dynamic Voronoi model” because the effective weight of each colony corresponds to its expansion rate and depends on the age of the colony, or equivalently on the distance from the inoculation centre, thereby introducing distance-dependent weights. In this approach, the petri dish was represented as a two-dimensional Cartesian grid, and each inoculation position **x***_i_* was treated as the origin of an expanding colony. For each colony, an experimentally fitted front propagation function *T_i_*(*d*) was defined, describing the time required for the colony edge to reach a radial distance *d* from its inoculation centre. This formulation allows the experimentally observed front dynamics to deviate from simple constant-speed linear growth and captures possible nonlinear acceleration or deceleration during expansion.

For any spatial position x on the grid, the Euclidean distance to inoculation centre *i* was first computed as

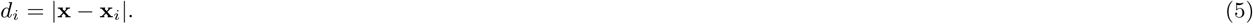

This geometric distance was then converted into an effective front arrival time

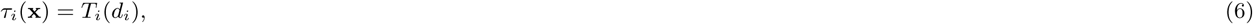

where *τ_i_*(x) denotes the predicted time at which colony *i* reaches position x according to its measured expansion law.

Each spatial grid point was subsequently assigned to the colony with the minimal arrival time,

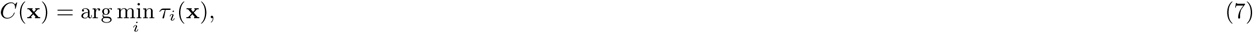

where *C*(x) denotes the colony index assigned to spatial position x in the predicted final partition, such that every location belongs to the colony whose expansion front is expected to arrive there before all competing colonies.

Repeating this minimization over the entire Petri dish yielded a weighted Voronoi tessellation in which competition boundaries correspond to equal front arrival times rather than equal geometric distances. In the general implementation, the front propagation functions *T_i_*(*d*) were represented by low-order polynomial fits obtained from the measured colony radius trajectories, thereby allowing the method to reproduce shifted, asymmetric, or curved interfaces generated by unequal colony expansion kinetics. Because this dynamic Voronoi model requires only experimentally measured front dynamics and negligible computational cost, it provides a practical quantitative framework for rapid forward design of target spatial patterns.

## Code availability

The code used to reproduce the reaction–diffusion simulations in one, two, and three dimensions presented in this study is available in the following GitHub repository: https://github.com/SchaerliLab/self_organized_voronoi.

## Acknowledgements

We thank Johannes Keegstra and Riccardo Foffi, ETH Zurich, for insightful discussions on bacterial motility and nutrient diffusion. This work was funded by the Swiss National Science Foundation (310030 200532 awarded to Y.S), a UNIL FBM PhD fellowship in Life Sciences (awarded to JH.P) and the University of Lausanne.

## Disclosure and competing interests statement

The authors declare no competing interests.

## Supplementary information

Supplementary Information. Supplementary Movies 1-3.

## Notes

### Competing Interest Statement

The authors have declared no competing interest.

### Summary of Updates

A formatting issue in the previous version caused part of the manuscript text to appear as part of Figure 2 caption. This has now been corrected. Additionally, some references to Fig. 4 actually referred to Fig. 3, and this has now also been corrected.

