## Supplementary Information for "Harnessing *Escherichia coli* motility to engineer bacterial Voronoi patterns"

### 1 Additional Experimental and Simulation Results

#### 1.1 The effect of agar concentration on gap formation in Voronoi patterns

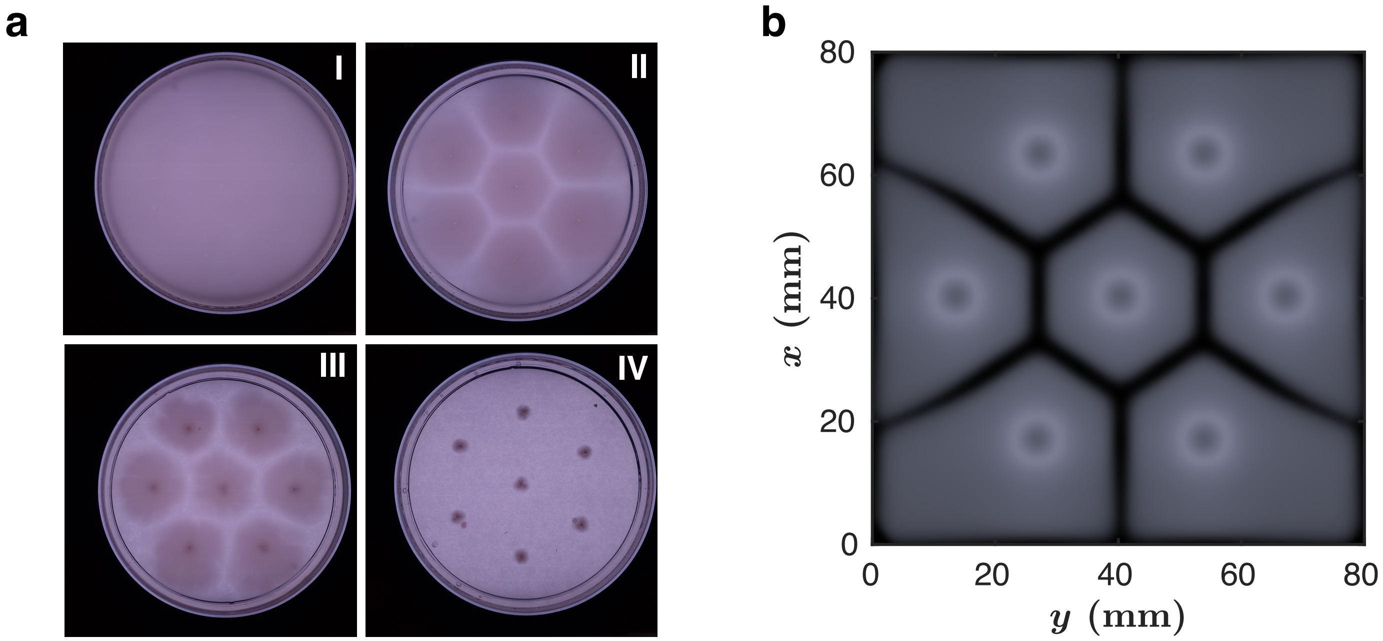

**Figure S1: Effect of agar concentration on bacterial expansion and gap formation.** (a) Expansion experiments with *E. coli* MG1655 cells in which seeds were inoculated at identical initial positions, with varying agar concentrations, I: 0.225%, II: 0.35%, III: 0.45%, IV: 0.50%. (b) Reaction–diffusion 2D model of bacterial expansion with parameters reflecting the 0.35% agar condition.

#### 1.2 Additional results of strain-dependent pattern formation

The Fisher-type reaction–diffusion model produces a constant expansion rate for each strain, corresponding to a fixed weight in the resulting weighted Voronoi pattern. In experiments, however, expansion rates can vary with time or position, leading to dynamically changing effective weights and, consequently, to curved or otherwise non-static Voronoi boundaries. To examine this effect within the reaction–diffusion model, we introduced a time-dependent diffusion coefficient for the blue strain, $D_{\mathrm{B}}(t) = D_{\mathrm{B},0}\,[1 + 2t^n/(t^n + T_{\mathrm{switch}}^n)]$, with $T_{\mathrm{switch}} = 20, \mathrm{h}$ and $n = 4$. This gradual increase in the effective cell diffusion, and hence expansion rate, produces time-dependent boundary curvature in the weighted Voronoi pattern (Figure S2e).

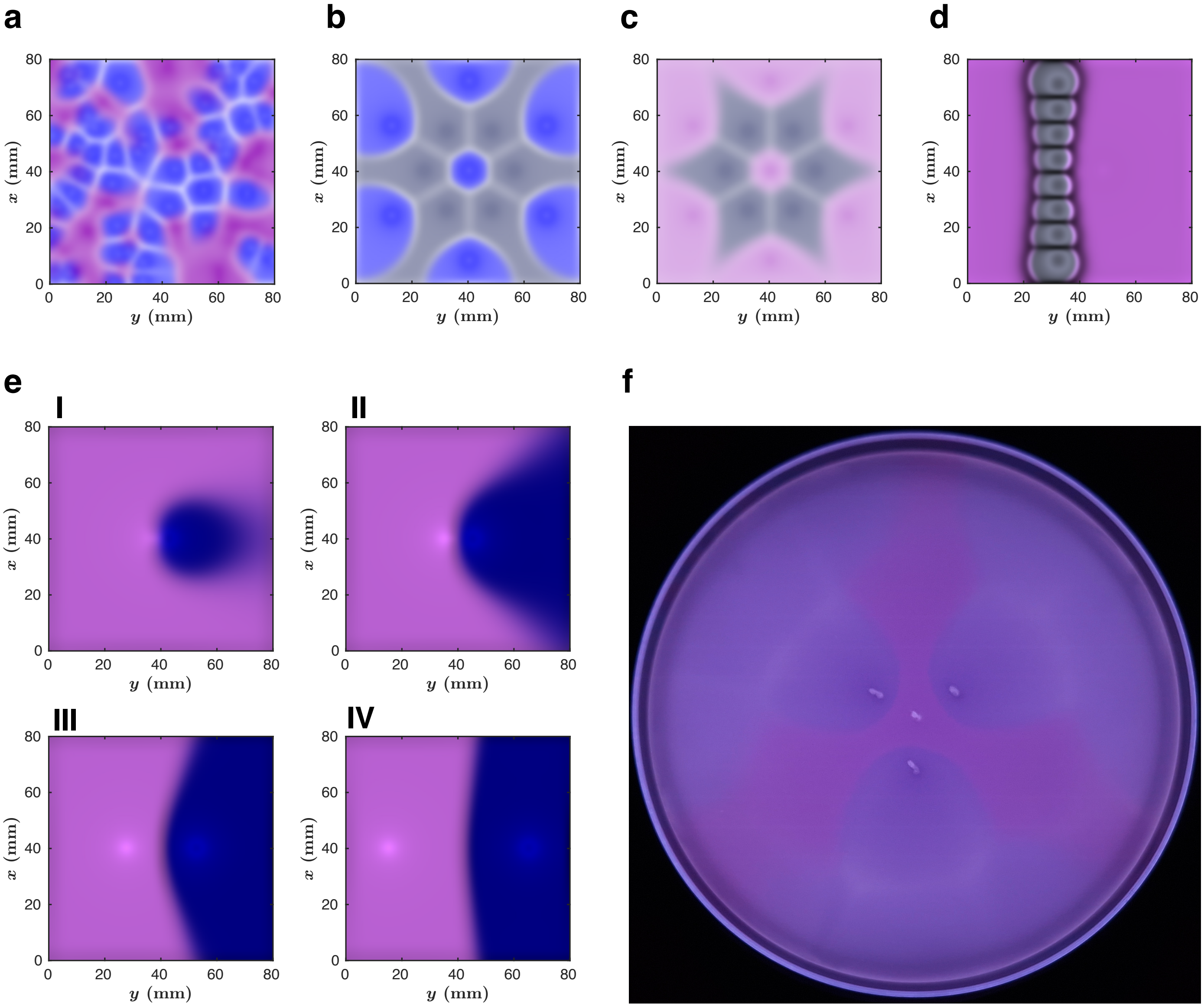

**Figure S2: Strain expansion rates and spatial arrangement determine Voronoi patterns.** Panels "a" to "e" represent the 2D reaction–diffusion model simulations of experiments from Figure 2c-g. (a) Simulation of a weighted Voronoi pattern with a randomly seeded configuration. Fast-expanding sectors (pink) grow around and constrain the slower-expanding sectors (blue). (b - c) Simulations showing that distinct spatial patterns can emerge from the same initial inoculation arrangement, by only varying the strain expansion rates. WT (gray), Amil-CP (blue), and TS-Purple (pink). (d) 2D model of an extremely slow-expanding strain (gray) inoculated in a barrier configuration, with a single inoculation of the fast-expanding TS-Purple strain (pink) on the right side. (e) 2D simulation showing how varying distances between the inoculated strains (pink - TS-Purple, blue - Amil-CP) affect the boundary shape (I - 5mm, II - 10mm, III - 25mm and IV - 50mm). (f) Experimental data showing unconventional patterns generated by TS-Purple and Amil-CP strains through distance-dependent expansion rates.

#### 2 The reaction–diffusion model

We consider the following reaction–diffusion equations describing the dynamics of the cell density $c_i$ and nutrient concentration $f$. The biological interpretation of the model is given in the main text; here we derive the dimensionless form of the governing equations.

$$\partial_t c_i = \nabla(D_{c,i}(f)\nabla c_i) + \frac{1}{2}\nabla(c_i \nabla D_{c,i}(f)) + k_i \frac{f}{f + K_{c,i}} c_i, \tag{S1}$$

$$\partial_t f = D_f \nabla^2 f - \sum_i \omega_i k_i \frac{f}{f + K_{c,i}} c_i, \tag{S2}$$

$$D_{c,i}(f) = D^0_{c,i} \frac{f}{f + K_f}. \tag{S3}$$

The quantities $c_i$ and $f$ cannot be measured on an absolute concentration scale in our experiments. Instead, the measured cell signal originates from light scattering and pigment absorption, while the nutrient consists of a complex LB medium with no directly measurable concentration. Consequently, we formulate the model using dimensionless concentrations. Besides simplifying the comparison with experiments, this transformation removes the nutrient conversion factor $\omega_i$, which represents the amount of nutrient required to produce one unit of biomass of cell type $i$.

Using the initial nutrient concentration $f_0$ as the characteristic concentration scale, we introduce the dimensionless variables

$$\hat{f} = \frac{f}{f_0}, \qquad \hat{c}_i = \frac{\omega_i c_i}{f_0}, \qquad \hat{K}_f = \frac{K_f}{f_0}, \qquad \hat{K}_{c,i} = \frac{K_{c,i}}{f_0}. \tag{S4}$$

Substituting these variables into the governing equations yields the dimensionless model presented in Equations 1–3 of the main text. For simplicity of notation, the hats are omitted throughout the remainder of the manuscript.

#### 3 (Anti-)Gap formation

##### 3.1 Travelling front profile

In the experiments, we observe expanding bacterial colonies that propagate as travelling fronts. Our observations indicate that slowly expanding colonies develop a depleted region when two fronts meet, corresponding to a negative surplus at the propagating front. In contrast, rapidly expanding colonies produce a positive surplus, resulting in a local accumulation of cells after the collision.

The numerical simulations reproduce this behaviour and converge to a travelling-wave solution. As shown in Figure S3, the cell concentration profile propagates from left to right, while nutrients are continuously converted into biomass (F $\rightarrow$ C), giving rise to a receding nutrient front. Because nutrient consumption is directly coupled to cell production, the sum of the cell and nutrient concentrations represents the total amount of biomass available. Remarkably, the quantity $c + f$ is not spatially uniform within the travelling wave. Instead, it exhibits either a local depletion or a local surplus at the propagating front (Figure S3, green curve). When two travelling fronts collide, this local imbalance is preserved and gives rise to either a depletion zone (gap) or an accumulation zone (antigap) between the colonies.

As shown in the following section, the sign of this surplus is determined solely by the relative transport of cells and nutrients. Specifically, when the effective cell diffusion coefficient is smaller than the nutrient diffusion coefficient ($D_c < D_f$), the propagating front develops a negative surplus, leading to a gap after collision. Conversely, when the effective cell diffusion exceeds nutrient diffusion ($D_c > D_f$), a positive surplus is formed, resulting in an antigap between the colonies.

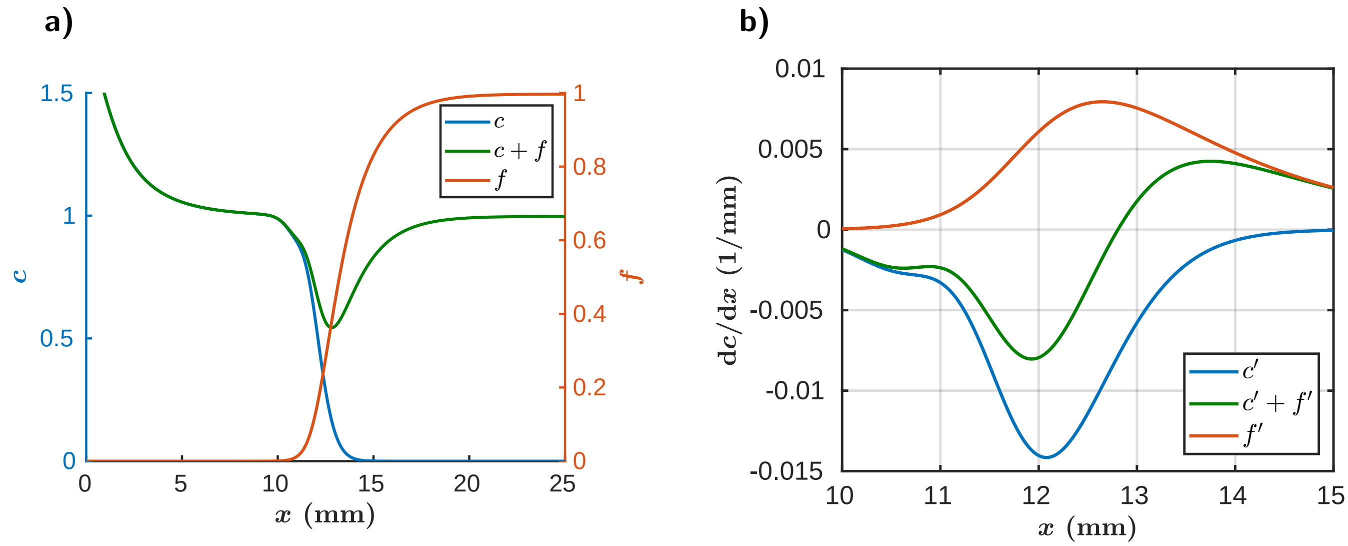

**Figure S3: Travelling-wave profiles.** **(a)** Concentration profiles of the cells $c$ (blue), nutrients $f$ (orange), and their sum $c + f$ (green) in the function of space $x$. The sum of the two concentrations exhibits a local negative surplus at the propagating front, when $D_c/D_f = 0.1$ **(b)** Spatial derivatives of the cell and nutrient concentration profiles, together with the derivative of their sum, shown using the same color scheme as in panel (a). The derivatives highlight the relative positions and widths of the sigmoid concentration profiles, which determine the magnitude and sign of the surplus.

#### 3.2 Estimating the surplus in the travelling front

To gain analytical insight into the origin of the cell surplus at the propagating front, we consider a simplified version of the reaction–diffusion model (Equations 1–2) for a single cell type with a constant cell diffusion coefficient. The model describes the dynamics of the cell density, $c(x, t)$, and nutrient concentration, $f(x, t)$, according to

$$\frac{\partial c}{\partial t} = D_c \nabla^2 c + R(f, c), \tag{S5}$$

$$\frac{\partial f}{\partial t} = D_f \nabla^2 f - R(f, c), \tag{S6}$$

where $D_c$ and $D_f$ denote the effective diffusion coefficients of cells and nutrients, respectively, and $R(f, c)$ is the local proliferation rate. We assume that the colony expands with a constant velocity $v$ and introduce the comoving coordinate $\xi = x - vt$, together with the stationary travelling-wave profiles $C(\xi) = c(x, t)$, $F(\xi) = f(x, t)$, where primes denote differentiation with respect to $\xi$. The governing equations become

$$-vC' = D_c C'' + R(F, C), \tag{S7}$$

$$-vF' = D_f F'' - R(F, C). \tag{S8}$$

Defining the total concentration

$$U(\xi) = C(\xi) + F(\xi), \tag{S9}$$

and adding the two equations eliminates the reaction term,

$$-vU' = D_c C'' + D_f F''. \tag{S10}$$

Using the identity $U'' = C'' + F''$ gives

$$-vU' - D_f U'' = (D_c - D_f)C''. \tag{S11}$$

Remarkably, the reaction term cancels exactly, implying that the surplus is determined solely by transport. For a steadily propagating travelling wave, the advection term dominates the diffusion of the total concentration, $|vU'| \gg |D_f U''|$, such that Equation S10 can be approximated by

$$-vU' \approx (D_c - D_f)C''. \tag{S12}$$

Integrating Equation S12 from $-\infty$ to $\xi$, and using the boundary conditions $C(-\infty) = 1$, $F(-\infty) = 0$, $C'(-\infty) = 0$, yields

$$-v(U - 1) = (D_c - D_f)C'. \tag{S13}$$

Defining the surplus

$$\delta U = U - 1, \tag{S14}$$

gives

$$\delta U = -\frac{D_c - D_f}{v} C'. \tag{S15}$$

Since $v > 0$ and $C' < 0$ throughout the travelling front, the surplus is positive for $D_c > D_f$ and negative for $D_c < D_f$. In Section 3.2.1 we give an alternative derivation for Equation S15.

Finally, we can also give an estimation for the peak height of the surplus. Assuming that the cell profile is well approximated by an error function,

$$C(\xi) = \frac{1}{2}\left(1 - \operatorname{erf}\left(\frac{\xi}{w}\right)\right), \tag{S16}$$

its derivative becomes

$$C'(\xi) = -\frac{1}{w\sqrt{\pi}} \exp\left(-\frac{\xi^2}{w^2}\right), \tag{S17}$$

where $w$ denotes the characteristic front width. The gradient reaches its maximum magnitude at the center of the front,

$$C'(0) = -\frac{1}{w\sqrt{\pi}}. \tag{S18}$$

Substituting this result into Equation S15 gives an estimate for the maximum surplus,

$$\delta U_{\max} = \frac{D_c - D_f}{vw\sqrt{\pi}}. \tag{S19}$$

Approximating the travelling wave by a Fisher–KPP front, $v = 2\sqrt{kD}$, $w \sim \sqrt{\frac{D}{k}}$, leads to

$$\delta U_{\max} \sim \frac{D_c - D_f}{D}, \tag{S20}$$

showing that the maximum surplus is primarily determined by the relative magnitudes of the cell and nutrient diffusion coefficients, while the dependence on the proliferation rate $k$ cancels to leading order. Consequently, a positive surplus is expected for $D_c > D_f$, whereas $D_c < D_f$ produces a depletion.

##### 3.2.1 Alternative derivation

An equivalent expression can be obtained by integrating Equation S10 directly,

$$-v(U-1) = D_c C' + D_f F'. \tag{S21}$$

Since nutrient consumption is locally coupled to biomass production, the two fronts approximately satisfy $F' \approx -C'$ within the reaction zone. Substituting this approximation immediately yields

$$-v(U-1) \approx (D_c - D_f)C', \tag{S22}$$

which is identical to Equation S15. Although this derivation relies on the additional approximation $F' \approx -C'$, it provides an intuitive interpretation: the surplus arises from the imbalance between the diffusive fluxes of cells and nutrients, while the reaction terms cancel through mass conservation.

### 4 Parameter estimation

#### 4.1 Diffusion coefficients and growth rates

The Fisher–KPP parameters reported in Table S1 were obtained by combining measurements from liquid growth assays and colony expansion experiments on agar plates. The cellular growth rate $k$ was determined from liquid culture growth curves by fitting only the initial exponential growth phase, before nutrient depletion became significant. To account for the short adaptation period after inoculation, we fitted the optical density measurements with an exponential growth model including a lag phase,

$$c(t) = c_0 \left[\exp(k(t - T_{\text{lag}})) \cdot \Theta(t - T_{\text{lag}}) + \Theta(T_{\text{lag}} - t)\right], \tag{S23}$$

where $c(t)$ is the measured optical density, $c_0$ is the initial optical density, $k$ is the exponential growth rate, $T_{\text{lag}}$ is the lag time, and $\Theta$ denotes the Heaviside step function. Before the onset of exponential growth ($t < T_{\text{lag}}$), the model predicts a constant optical density $c_0$, whereas after the lag phase the population grows exponentially with rate $k$. The model parameters were estimated by nonlinear least-squares fitting using MATLAB's `fit` function (Table S1).

The colony expansion velocity $v$ was determined independently from time-lapse measurements of single colonies growing on agar plates. The colony radius was measured as a function of time, and a linear function was fitted to the initial region where the colony expanded with an approximately constant velocity. The slope of this fit yielded the expansion velocity $v$. Finally, the effective diffusion coefficient was calculated from the Fisher–KPP relation $v = 2\sqrt{kD}$, which gives $D = \frac{v^2}{4k}$. Confidence intervals for $k$ and $v$ were obtained directly from the nonlinear and linear fits, respectively. The corresponding confidence interval for the effective diffusion coefficient $D$ was estimated by propagating these uncertainties through the Fisher–KPP relation.

#### 4.2 Front speed estimation

To verify that the experimentally estimated parameters are consistent with the Fisher–KPP prediction for front propagation, we performed numerical simulations of a single colony expanding in a radially symmetric (polar) coordinate system

**Table S1:** Estimated Fisher–KPP parameters for the different experimental conditions. Values are reported as estimate ± 95% confidence interval from the fitted curve. The expansion velocity $v$ was obtained from colony expansion measurements, the growth rate $k$ from growth curve analysis, and the effective diffusion coefficient $D$ was calculated from the Fisher–KPP relation $v = 2\sqrt{kD}$. For conditions lacking growth curve measurements, only the expansion velocity $v$ could be determined, while $k$ and $D$ are not available. Low density is LB 0.225% agar, and high density is LB 0.35% agar.

| Strain and Condition | $v$ (mm h$^{-1}$) | $k$ (h$^{-1}$) | $D$ (mm$^2$ h$^{-1}$) |
|---|---|---|---|
| MG1655 WT (low density) | $4.207 \pm 0.123$ | $1.387 \pm 0.094$ | $3.191 \pm 0.285$ |
| MG1655 WT (high density) | $1.664 \pm 0.015$ | $1.387 \pm 0.094$ | $0.499 \pm 0.035$ |
| AHL-induc. cheZ (low density) 0 nM AHL | $1.617 \pm 0.106$ | $1.449 \pm 0.041$ | $0.484 \pm 0.065$ |
| AHL-induc. cheZ (low density) 0.2 nM AHL | $2.204 \pm 0.127$ | $1.445 \pm 0.045$ | $0.900 \pm 0.107$ |
| AHL-induc. cheZ (low density) 2 nM AHL | $0.861 \pm 0.083$ | $1.456 \pm 0.043$ | $0.137 \pm 0.027$ |
| AHL-induc. cheZ (low density) 10 nM AHL | $0.641 \pm 0.094$ | $1.350 \pm 0.042$ | $0.076 \pm 0.023$ |
| AHL-induc. cheZ (high density) 0 nM AHL | $0.955 \pm 0.021$ | $1.449 \pm 0.041$ | $0.157 \pm 0.008$ |
| AHL-induc. cheZ (high density) 0.2 nM AHL | $0.940 \pm 0.025$ | $1.445 \pm 0.045$ | $0.153 \pm 0.009$ |
| AHL-induc. cheZ (high density) 2 nM AHL | $0.627 \pm 0.024$ | $1.456 \pm 0.043$ | $0.067 \pm 0.006$ |
| AHL-induc. cheZ (high density) 10 nM AHL | $0.218 \pm 0.015$ | $1.350 \pm 0.042$ | $0.009 \pm 0.001$ |
| AHL Sender (low density) | $2.395 \pm 0.129$ | $1.238 \pm 0.040$ | $1.158 \pm 0.130$ |
| Amil-CP (low density) | $1.897 \pm 0.145$ | $1.249 \pm 0.055$ | $0.720 \pm 0.114$ |
| TS-Purple (low density) | $3.964 \pm 0.110$ | $1.377 \pm 0.024$ | $2.853 \pm 0.166$ |
| Amil-CP (high density) | $0.978 \pm 0.033$ | $1.249 \pm 0.055$ | $0.192 \pm 0.016$ |
| TS-Purple (high density) | $1.844 \pm 0.100$ | $1.377 \pm 0.024$ | $0.618 \pm 0.068$ |
| MG1655 ΔcheBΔcheR (low density) | $0.430 \pm 0.010$ | $1.616 \pm 0.131$ | $0.029 \pm 0.003$ |
| L-arab-induc. cheZ (high density), 0% arab. | $0.118 \pm 0.004$ | $1.878 \pm 0.134$ | $0.002 \pm 0.0002$ |
| L-arab-induc. cheZ (high density), 0.0001% arab. | $0.577 \pm 0.017$ | $1.9831 \pm 0.132$ | $0.042 \pm 0.004$ |
| L-arab-induc. cheZ (high density), 0.0005% arab. | $1.061 \pm 0.017$ | $1.933 \pm 0.164$ | $0.146 \pm 0.013$ |
| L-arab-induc. cheZ (high density), 0.02% arab. | $1.163 \pm 0.043$ | $1.722 \pm 0.168$ | $0.196 \pm 0.024$ |

(Fig. SS4). The model describes the dynamics of the cell density $c(r,t)$ and nutrient concentration $F(r,t)$ according to

$$\frac{\partial c}{\partial t} = D\nabla^2 c + kcF, \tag{S24}$$

$$\frac{\partial F}{\partial t} = D_F\nabla^2 F - kcF, \tag{S25}$$

where $D$ is the experimentally estimated effective cell diffusion coefficient, $D_F$ is the nutrient diffusion coefficient [1], and $k$ is the growth rate measured from liquid growth curves. Simulations were performed on a one-dimensional radial domain of radius $R = 50$ mm using 1001 spatial grid points and explicit finite differences. Initially, cells occupied a circular colony of radius $R_0 = 2$ mm with density $c = 1$, while nutrients were depleted within the inoculated region and set to $F = 1$ elsewhere. On the boundary no flux boundary condition was used. The front position was defined as the radial location where the cell density exceeded a threshold value of $c = 0.1$. The resulting front trajectory was fitted by a linear function to obtain the asymptotic propagation speed $v$. Figure S4a shows representative cell and nutrient concentration profiles during colony expansion, while Fig. S4b shows the corresponding front position as a function of time together with the fitted linear trend. The measured front speed was in good agreement with the Fisher–KPP prediction $v = 2\sqrt{kD}$, demonstrating that the experimentally determined growth and diffusion parameters accurately capture the observed expansion dynamics.

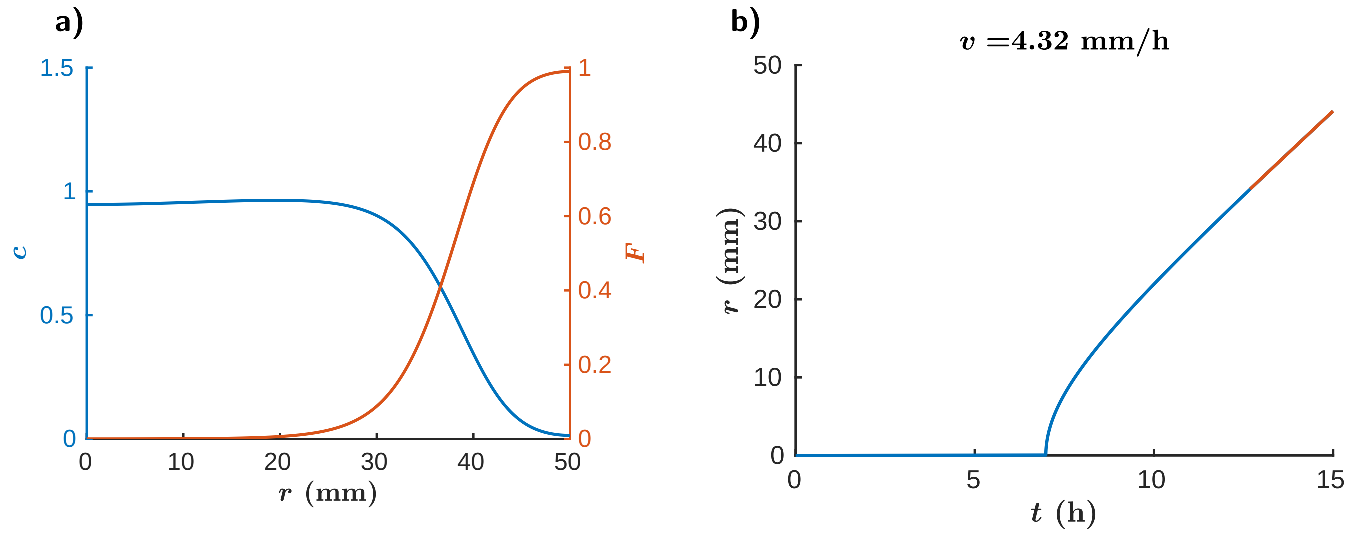

**Figure S4: Validation of the Fisher–KPP front speed prediction in a radially expanding colony. (a)** Simulated cell density ($c$) and nutrient concentration ($F$) profiles during colony expansion in a one-dimensional polar coordinate system at the end of the simulation ($T = 15$ hours). Cells proliferate by consuming nutrients and spread through diffusion, resulting in the formation of a traveling front. **(b)** Front position as a function of time, determined from the radial location where the cell density exceeds a threshold value ($c = 0.1$). The solid red line shows a linear fit to the asymptotic front trajectory, yielding the propagation speed $v$. The simulated front speed is consistent with the Fisher–KPP prediction $v = 2\sqrt{kD}$ based on the experimentally measured growth rate $k$ and effective diffusion coefficient $D$ tested with the WT (low density) strain (Table S1). The experimentally measured front speed was 4.21 mm/h, compared with 4.32 mm/h obtained from the simulation.

#### 4.3 Half-saturation constant in the Monod function

In the reaction–diffusion model, cellular proliferation is described by a Monod function (Equation 1). As discussed in Section 4.1, the maximal growth rate $k_i$ was obtained by fitting the initial exponential phase of the liquid culture growth curves. During this early phase, nutrient concentrations remain high and the Monod function is close to its maximal value, making the estimate of $k_i$ largely independent of the half-saturation constant $K_{c,i}$. The half-saturation constant becomes more important once nutrients become limiting. In particular, when two strains occupy the same region of the agar, they compete for the same nutrient pool, and $K_{c,i}$ determines how efficiently each strain can continue growing at low nutrient concentrations. Consequently, this parameter influences the competitive outcome even if the maximal growth rates are identical (Figure 4).

The Monod function used in the reaction–diffusion model is intentionally a simplified description of nutrient-limited growth. Rich LB medium contains numerous carbon and nitrogen sources that are consumed sequentially or simultaneously, producing the characteristic multi-phase growth curves frequently observed in *E. coli*. To capture this behaviour, we fitted the experimental liquid culture measurements using a model with two independent nutrient pools,

$$\frac{\mathrm{d}c}{\mathrm{d}t} = \left( k_1 \frac{f_1}{f_1 + K_1} + k_2 \frac{f_2}{f_2 + K_2} \right) c, \tag{S26}$$

$$\frac{\mathrm{d}f_1}{\mathrm{d}t} = -\omega_1 k_1 \frac{f_1}{f_1 + K_1} c, \tag{S27}$$

$$\frac{\mathrm{d}f_2}{\mathrm{d}t} = -\omega_2 k_2 \frac{f_2}{f_2 + K_2} c, \tag{S28}$$

where $c$ is the optical density of the culture, $f_1$ and $f_2$ denote the normalized concentrations of a rapidly and a slowly consumed nutrient pool, respectively, $k_i$ are the maximal growth rates, $K_i$ are the corresponding Monod half-saturation constants, and $\omega_i$ describe the nutrient consumption required to produce one unit of biomass.

The model parameters were estimated by nonlinear least-squares fitting using MATLAB's `lsqnonlin` function (Figure S5). For every parameter set, the ordinary differential equations were solved numerically using `ode45`, and the simulated optical density was compared with the experimental growth curve. The optimization minimized the sum of squared residuals between the measured and simulated optical densities.

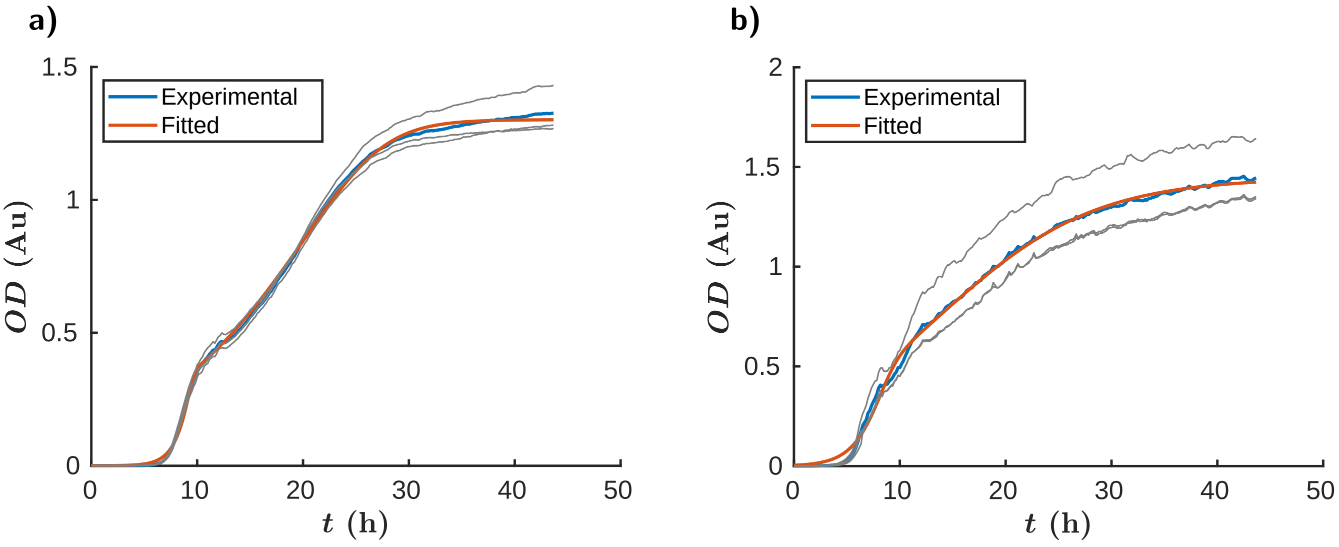

**Figure S5: Fits to the experimental growth curves.** Blue curves show the experimental measurements (mean of three biological replicates). Grey curves show the individual biological replicates. Red curves show the corresponding fits of the two-nutrient Monod model. **(a)** Amil-CP strain. **(b)** TS-Purple strain.

The primary purpose of fitting the two-nutrient model was not to determine all model parameters, but to estimate the relative magnitudes of the Monod half-saturation constants for the different strains. The fitted values indicate that the Amil-CP strain can continue growing efficiently at substantially lower nutrient concentrations than the TS-Purple strain. For the dominant nutrient pool, we obtained $K_1 = 0.21$ for the Amil-CP strain and $K_1 = 0.94$ for the TS-Purple strain. These values support the parameter choice used in the reaction–diffusion simulations of mixed colonies, where we set $K_c = 0.1$ for the Amil-CP strain and $K_c = 0.5$ for the TS-Purple strain. Since the reaction–diffusion model is phenomenological and represents all nutrients by a single effective nutrient concentration, these values should be interpreted as effective half-saturation constants rather than direct biochemical parameters. In all other simulations, we used $K_c = 0.1$. The precise value of this parameter does not qualitatively affect the resulting patterns, as long as a single strain occupies the colony. It becomes important only when multiple strains compete locally for the same nutrient pool, where differences in the half-saturation constants influence their competitive advantage under nutrient-limited conditions.

### 5 Strains, plasmids and primers list

| Strain | Genomic modifications | Plasmid(s) | Figures |
|---|---|---|---|
| MG1655 Wild-type | | | 1a-dI-dII, 2a-d-e, 3b |
| MG1655 ΔcheZ | cheZ deletion | | |
| MG1655 ΔcheBΔcheR | CheB and CheR deletions | | 1dIII-dIV-g, 2gII-gIII |
| Amil-CP (Blue) | | amilCP | 2a-c-d-f, 4a |
| TS-purple (Pink) | | tsPurple | 2a-c-e-f-g, 4a |
| L-arab-induc. cheZ | cheZ deletion | pBAD-cheZ | 3b-c |
| AHL-induc. cheZ (Receiver) | cheZ deletion | pLux-cheZ-mCherry + pLuxR-LasI | 3e-f, 4c-d |
| AHL sender | | Toggle Switch - LuxI + tsPurple | 4c-d |

**Table 1: Strains used in this work.**

| Plasmid | Features | Resistance | Origin | Source |
|---|---|---|---|---|
| amilCP | amilCP chromoprotein | Chloramphenicol | pSB1C3 | Addgene ID #117847 |
| tsPurple | tsPurple chromoprotein | Chloramphenicol | pSB1C3 | Addgene ID #117848 |
| pBAD-cheZ | Constitutive expression of the AraC regulator, arabinose-inducible expression of CheZ via the pBAD promoter | Spectinomycin | pCDF | This work |
| pLux-cheZ-mCherry | Constitutive expression of the mCherry reporter, AHL-inducible expression of CheZ via the pLux promoter | Kanamycin | p15A | This work |
| pLuxR-LasI | Constitutive expression of the quorum sensing receptor LuxR and the synthase LasI | Chloramphenicol | pMB1 | Described in [2] |
| Toggle Switch - LuxI | inducible *luxI* production | Kanamycin | pColE1 | Addgene ID #251147 |

**Table 2: Plasmids used in this work.**

| Primer | Sequence | Function |
|---|---|---|
| PR-EB-196 | ttatcagaccgcctgatatgacgtggtcac gccacatcaggcaatacaaagtgtaggct ggagctgcttc | Amplification of the FRT-KamR-FRT cassette with homology for the *cheZ* genomic region |
| PR-EB-197 | ggaaaaactcaacaaaatctttgagaaact gggcatgtgaggatgcgactattccgggga tccgtcgacc | Amplification of the FRT-KamR-FRT cassette with homology for the *cheZ* genomic region |
| PR-EB-198 | gagtaaggggcaaaacaggc | Colony PCR of the *cheZ* genomic region |
| PR-EB-199 | gctgaaaacaattcgtgcgg | Colony PCR of the *cheZ* genomic region |
| PR-EB-239 | tcaaaatccaagactatccaacaaatcg | Amplification of genomic *cheZ* |
| PR-EB-240 | atgatgcaaccatcaatcaaacc | Amplification of genomic *cheZ* |
| PR-EB-243 | gacgatttgttggatagtcttggattttga aagagatttctacacgattgagcac | Cloning of *cheZ* in a pCDF vector |
| PR-EB-245 | ttgattgatggttgcatcatcggcgcctcc ctatcagt | Cloning of *cheZ* in a pCDF vector |
| PR-EB-161 | tcgctgggacgcccgagtattaactatcgt tcaactg | Gibson assembly of pBAD-cheZ, amplifies the backbone |
| PR-EB-193 | gccgcaactagtacaatcgc | Gibson assembly of pBAD-cheZ, amplifies the backbone |
| PR-EB-272 | gcgattgtactagttgcggcagagtcttga agtggtggcc | Gibson assembly of pBAD-cheZ, amplifies AraC and pBAD |
| PR-EB-273 | atactcgggcgtcccagcgaggacggtaaa gtttccagaatctatcc | Gibson assembly of pBAD-cheZ, amplifies AraC and pBAD |
| PR-EB-341 | atgatgcaaccatcaatcaaacctgc | Addition of an ATG codon to the *cheZ* sequence in the original version of pLux-cheZ-mCherry |
| PR-EB-342 | gcaggtttgattgatggttgcatcatggtt tcctgtgtgagaattctttattcg | Addition of an ATG codon to the *cheZ* sequence in the original version of pLux-cheZ-mCherry |
| PR-EB-006 | tcaagttggacatcacctccc | Gibson assembly of pLux-cheZ-mCherry, binds inside the mCherry sequence |
| PR-EB-150 | cgcgttcgtactgttccac | Gibson assembly of pLux-cheZ-mCherry, binds inside the mCherry sequence |

**Table 3: Primers used in this work.**
